# Characterization of the maize (*Zea mays*)-*Ustilago maydis* interaction in a warming climate

**DOI:** 10.1101/2025.05.26.656155

**Authors:** Christian G. Schwarz, Finn Hartmann, Christopher Zier, Karina van der Linde

## Abstract

Maize is among the most grown crops worldwide. Concurrently, infections with *Ustilago maydis*, causing corn smut disease, occur in all major growing regions. This is accompanied by loss of biomass, yields, and silage quality. As global warming increases, modeling analysis predicts an increase in pathogen infestation. Nevertheless, there are no maize lines resistant to *U. maydis* and no effective fungicides available.

Based on the evaluation of climate data for the State of Bavaria (Germany), maize-*U. maydis* infection trials and RNAseq analysis were conducted on a variety of maize cultivars under different temperature conditions generating a large phenotypic and transcriptomic dataset to determine the influence of temperature changes and differences in plant susceptibility. Even a minimal increase in temperature resulted in increased symptoms and significant variances in expression. Infection-gene expression association analysis followed by *in vivo* tests identified GIBBERELLIC ACID STIMULATED TRANSCRIPT-LIKE4 (GSL4) and *γ*-aminobutyric acid as important factors for the infection.

## Introduction

Maize is among the most grown staple crops worldwide. Besides this it is a major source of renewable energy. In Germany, the State of Bavaria is the biggest maize producer ^1^. The biotrophic fungus *Ustilago maydis* is the causative agent of corn smut disease and can be found worldwide. *U. maydis* infects all aerial parts of maize resulting in tumor formation ^2^. This is associated with reduced plant biomass production and a deterioration in the maize silage properties ^3,4^.

Infection starts when two compatible haploid sporidia mate to form a dikaryotic filament on the plant surface. On the tip of the filament an appressorium is developed that penetrates the plant surface and the plant plasma membrane invaginates resulting in the establishment of a biotrophic interaction zone between the fungal hyphae and the plant cell ^5–7^. In the plant, *U. maydis* proliferates and branches while also inducing hyperplasic and hypertrophic plant tumor cell growth. Within these tumors, millions of fungal teliospores are produced. Initially, infection with *U. maydis* results in an unspecific plant defense response; once the biotrophic interaction is established, this defense is suppressed, tumor growth is initiated, and plant nutrients are diverted to the pathogen ^5,8,9^. Even though it is predicted that the genome of *U. maydis* encodes for 533 secreted effectors, no gene-for-gene resistance has been found ^10^. Instead, studies imply that infection frequency and severity is a polygenic trait ^11,12^. This is supported by data from infection studies of 28 maize cultivars where resistance levels ranged from tumor levels from 35 % to 94 % ^13^. Subsequently, multiple dQTLs (disease Quantitative Trait Loci) were identified, although the number varies significantly among existing studies ^14–16^. Despite the fact that the maize-*U. maydis* system serves as a model for biotrophic plant-pathogen interactions in research, little is known about the underlying mechanisms of plant susceptibility and its cultivar specific variation.

Climate change is defined as a lasting alteration of meteorological variables over an extended period of time. Today’s climate change is mostly a result of increased greenhouse gases, primarily from burning of fossil fuels, industrial processes, or deforestation ^17^. Greenhouse gases in the atmosphere absorb radiation energy emitted by earth and result in global warming through the greenhouse effect. Global warming involves both a steady temperature increase and induces a multitude of climate disturbances, e.g. changes in precipitation patterns, intensified droughts, heatwaves, and storms, impacting agriculture, water resources, and public health ^17,18^. Representative Concentration Pathways (RCPs) are scenarios used in prediction analysis to project future climate change. RCPs represent different trajectories of radiative forcing, which is a measure of the imbalance between incoming and outgoing energy in the Earth’s atmosphere ^19^. These pathways are characterized by varying levels of greenhouse gas emissions and are used to assess the potential impacts of different emission mitigation strategies on future climate conditions. For instance, RCP2.6 represents a stringent scenario, aiming to limit global warming to below 2 °C above pre-industrial levels ^20^. In contrast, RCP8.5 represents a high-emission pathway, projecting significant climate impacts by the end of the century ^21^.

Between 1980 and 2008, maize yields already have declined by 3.8% due to rising temperatures, and it is predicted that yields will decline by a further 10-22% until 2100 ^22,23^. Besides the direct impact of rising temperatures on maize yield, rising temperatures also promote pathogen infection. Increasing temperatures are predicted to boost the susceptibility of agricultural crops to pathogens and to increase the virulence of pathogens ^24,25^. In addition, weather extremes will increase ^26^. Yet, it remains unclear how *U. maydis* infection frequency and severity is impacted by temperature changes and what the underlying molecular mechanisms could be.

In this study, we used 17 EU NAM founders, which is a selected set of European maize inbreed cultivars representing large genetic diversity, plus the US inbreed cultivar B73 for infection studies under four different temperature conditions ^27^. These temperature conditions were chosen based on previous studies, but also based on modelled climate data that specifically represents the temperatures in May in Bavaria for the years 1985 and 2050. Additionally, we simulated a short heatwave of three days. Phenotypic data were collected on tumor formation, as well as leaf length and time point of melanized spore formation. Nine out of the 18 cultivars were subjected to transcriptomic profiling under all temperature conditions with or without *U. maydis* infection. These RNAseq data together with the phenotypic data were used to identify two previously undescribed plant factors for the *U. maydis*-maize interaction.

## Results

### Small temperature changes impact *U. maydis* symptoms

In the German State of Bavaria, maize is typically sown at the end of April up to the beginning of May. Subsequently, the seedling stage, which is mostly used in other studies, is reached in May. Climate data for Bavaria for the month of May from the Bavarian Environment Agency were used as a basis to model future temperatures according to three different climate change scenarios (RCP 2.6, RCP 4.5, or RCP 8.5) and asses current temperatures ^19,28^. These models describe the change of monthly maximum and minimum temperatures of the 30-year rolling average from 1985 to 2085 compared to the 30-year average of the reference period (1971-2000; hereafter called “1985”) ^28^. 1985 was selected as reference because it included the last month when global temperatures were below the 20th century average ^29^. Thus, the temperature conditions in May 1985, with a mean daytime temperature of 17.9 °C and a mean nighttime temperature of 6.9 °C, were used as reference conditions before the occurrence of climate change-induced temperature changes for all further experiments ^28^. For May 2050, representing the 30-year average from 2036-2065, the RCP4.5 scenario predicted an increase of the mean maximum temperature by 1.2°C and the mean minimum temperature by 1.5°C compared to 1985. Therefore, for May 2050 the predicted average daytime maximum temperature is 19.1°C and the average nighttime minimum temperature is 8.4°C. Besides a continuous small change in temperature, extreme weather events such as heatwaves are also predicted to occur more frequently due to climate change. Already in May 2021 a short-term maximum daily temperature of 31.3 °C has been recorded in Germany. Based on these data and previously reported studies, four different temperature conditions were established for *U. maydis*-maize infection experiments (Fig. 1A): 1. Standard (28 °C day, 25 °C night), 2. 1985 (17.9 °C day, 6.9 °C night); 3. 2050 (19.1 °C day, 8.4 °C night); or 4. Heatwave (3 days 35 °C day, 25 °C night, followed by 17.9 °C day, 6.9 °C night).

**Fig. 1:**
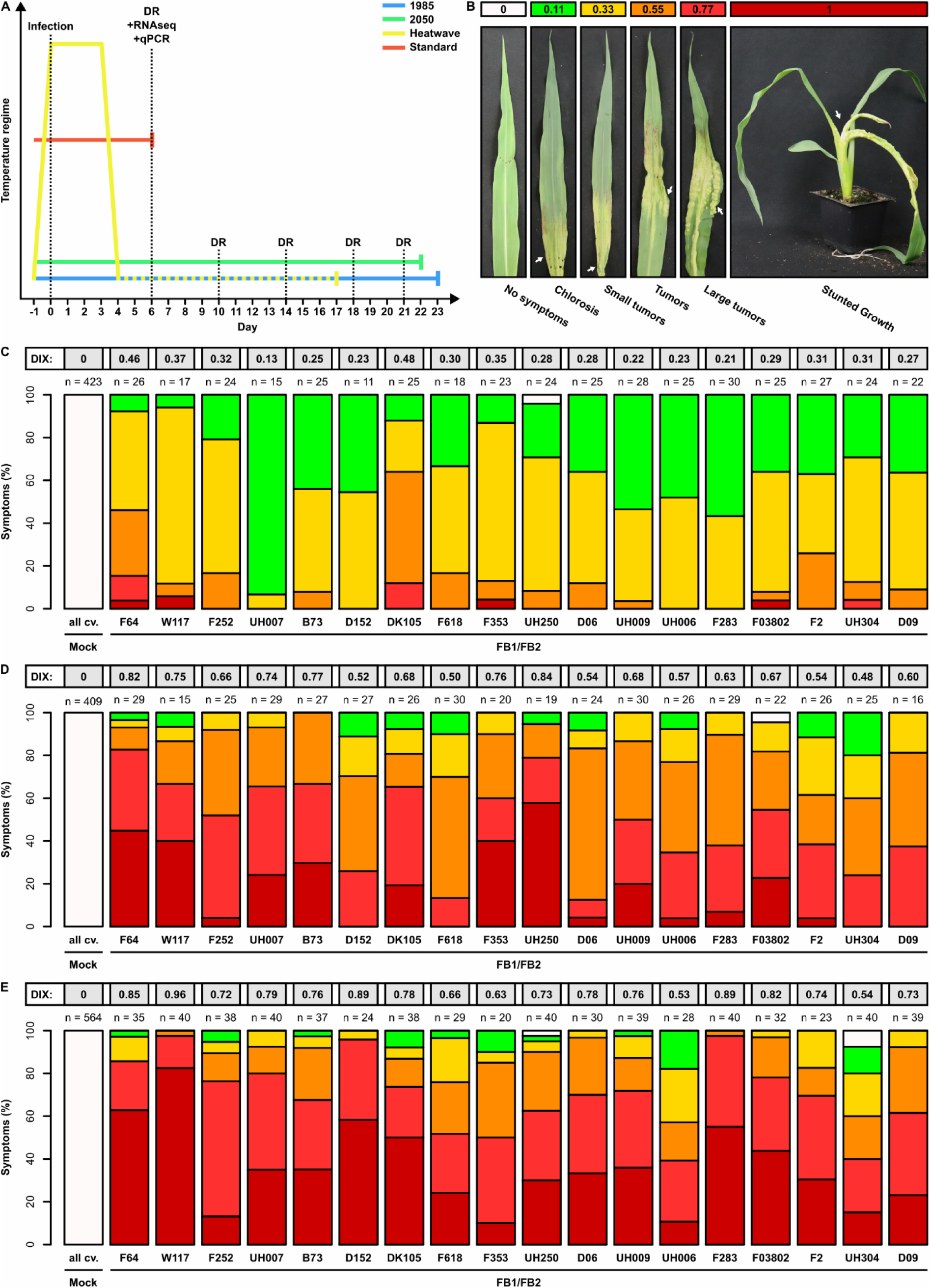
Comparative *U. maydis* infections of 18 maize cultivars under different temperature conditions. **A** Infection studies were performed under four different temperature regimes. **B** Photos of different infection symptoms. The different symptom categories are represented by specific colors and a specific disease index (DIX). **C-E** Disease ratings of 17 EU-NAM founder cultivars and B73 14 days after infection with the compatible *U. maydis* strains FB1 and FB2 or mock under the temperature conditions of 1985 (**C**), 2050 (**D**), or the temperature for 1985 with additional heatwave (**E**). All experiments were performed in two biologically independent replicates; the total number of plants used (n) is indicated in each case. The mean infection intensity for each maize cultivar is given as disease index (DIX).

Using temperatures for 1985, only chlorosis and small tumors were commonly observed 14 days after infection (Fig. 1C). Only in rare cases, large tumors or stunted growth occur in cultivars F64, W117, F353, or F03802. In contrast, under the slightly higher temperature conditions predicted for 2050, these severe infection symptoms occurred much more frequently, and were more severe overall (Fig. 1D). Stunted growth was found in 14 of the 18 tested cultivars. After the heatwave experiments, symptoms became more severe in most lines, with an increased incidence of large tumors and stunted growth (Fig. 1E). Quantitative PCR (qPCR) experiments were performed to determine fungal biomass in infected tissues, and these confirmed that the disease rating data reflected by increasing fungal presence. Besides disease ratings, the length of the fourth seedling leaf was measured after infection as an indicator of plant biomass production and compared to the leaf length of non-infected plants under the respective temperature conditions. Under all temperature conditions, leaf length was reduced by 12-27% as early as six days after infection with *U. maydis* in all maize cultivars. While the leaf length reduction under temperature conditions 1985 and 2050 continued to increase during the course of the experiment, no further significant leaf length reduction was measured under heatwave conditions later than 6 dpi. The greatest reduction (27%) was observed 14 days after infection under 2050 temperature conditions. Furthermore, spore formation was examined under all temperature conditions. While the first spores were formed on average 20-21 days after infection under the temperature conditions of 1985, sporulation occurred earlier both at the temperatures predicted for the year 2050 or after short-term heat stress. Under heatwave conditions, sporulation was observed in maize cultivars F2, F283, UH250, DK105, DK152, F252, W117, or F64 as early as 14 days after infection. Overall, disease progression as well as plant growth is accelerated under climate change conditions and sporulation appears to occur earlier at elevated temperatures in all cultivars. Additionally, large variances in response patterns to the different temperature conditions were observed between the used diverse cultivars for all phenotypic parameters.

### Transcriptomic variations in maize cultivars in response to different temperature conditions

To gain a deeper understanding of the molecular basis underlying the variance of phenotypic parameters between maize cultivars, transcriptomic analysis of nine maize cultivars (F64, F03802, W117, D06, D152, UH006, D09, B73, and UH250) was performed for mock treatments and *U. maydis* infections for all four temperature scenarios. B73 was included in RNAseq analysis as a representative of the US NAM panel. The EU NAM founder cultivars were chosen for RNAseq based on their response to *U. maydis* infection under different temperature conditions measured by disease ratings. The selected cultivars can be grouped into four different response categories: 1. Strong symptoms in all conditions (F64); 2. Low susceptibility in all conditions (UH006); 3. Average symptoms in all conditions (F03802, D006); 4. Fluctuating symptom strength compared to the other cultivars in different conditions (W117, D09, D152, UH250).

The RNAseq samples clustered in principal component (PC) 1 into a mock infected group or an *U. maydis* infected group (Fig. 2A). PC2 shows clustering of samples based on the temperature condition. A total of 33,059 DEGs (differentially expressed genes) were detected for all temperature conditions including 13,790 unique DEGs. 3,547 DEGs are shared among all conditions when comparing infected to mock-infected samples, while 156, 948, 667, 2,607 were specific to a certain temperature regiment (1985, 2050, Heatwave, or Standard respectively) (Fig. 2B). DEGs can be further divided into three categories for each temperature condition: 1. genes that show a change in expression in all nine maize lines examined (core DEGs); 2. DEGs that were identified in at least two maize lines (shared DEGs); 3. DEGs that were found in only one maize line (line-specific DEGs). Under 1985 temperature conditions, most DEGs were line-specific with few and core DEGs, whereas under standard temperature conditions the proportion of line-specific DEGs decreases and the proportion of core DEGs strongly increases (Fig. 2C).

**Fig. 2:**
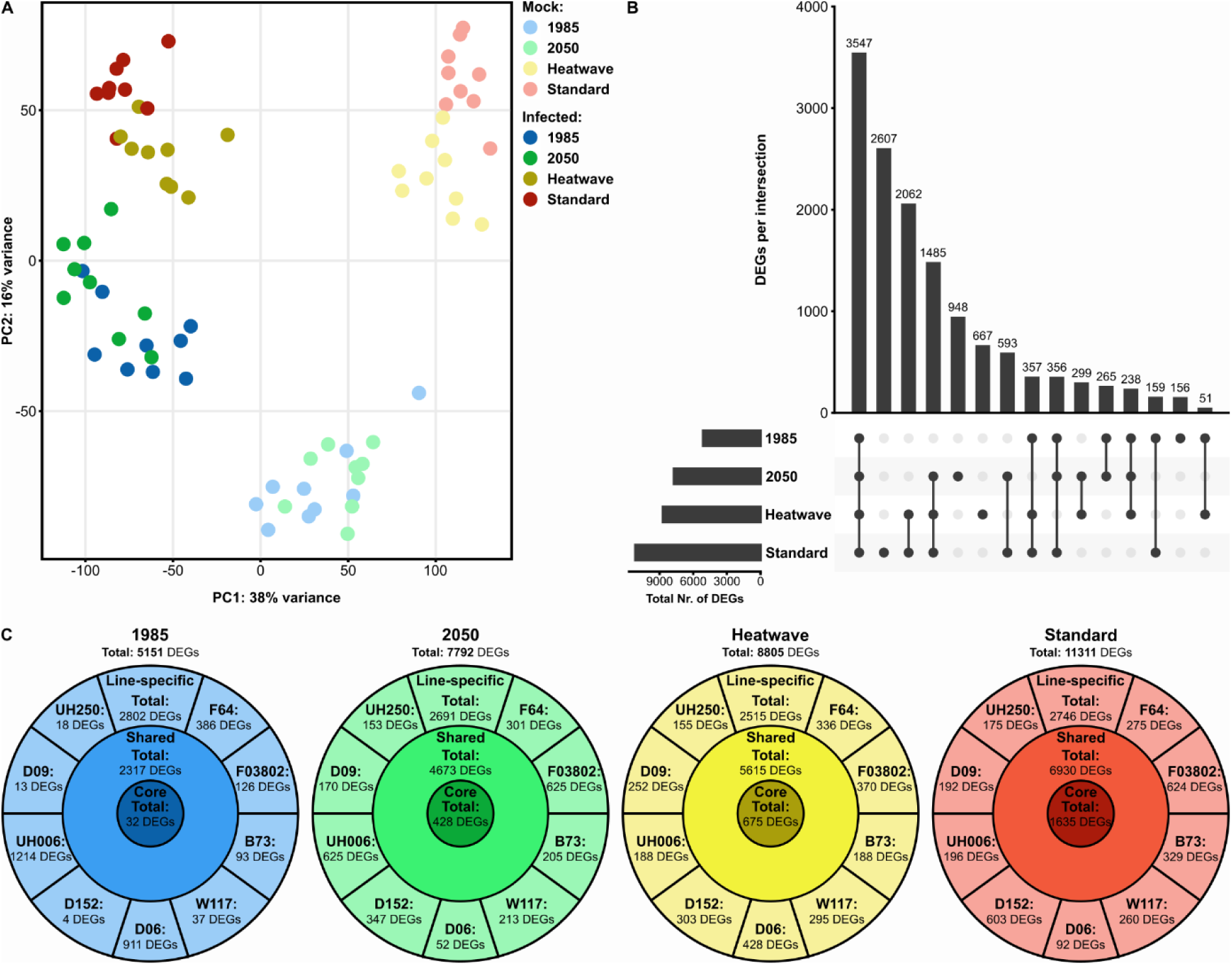
Gene expression analysis after *U. maydis* infection of maize cultivars at different temperatures. **A** Principal component analysis of the expression patterns of *U. maydis* infected and mock-infected samples under different temperature conditions: temperature conditions of 1985, 2050, the temperature for 1985 with additional heatwave, or standard conditions. Each dot represents RNAseq data of one of nine maize cultivars (B73, F03802, F64, D06, D09, D152, UH006, UH250, W117). **B** UpSet plot of differentially expressed maize genes (DEGs) under different temperature conditions. Genes with log_2_(expression fold change) > 2 or < 2 and adjusted *P* value < 0.001 were considered differentially expressed. For each temperature scenario the total number of DEGs is given. Only intersections including at least 50 genes are shown. Connecting black dots indicate overlap between conditions, while vertical bars represent the number of DEGs per intersection. **C** Distribution of DEGs at the different temperature conditions (temperature conditions of 1985 (blue), 2050 (green), the temperature for 1985 with additional heatwave (yellow), or standard condition (red)) in three categories: 1. Genes showing a significant change in expression in all nine maize cultivars tested (core DEGs); 2. DEGs identified in at least two maize cultivars (shared DEGs); 3. DEGs found in only one maize cultivar (line-specific DEGs).

To identify common patterns, which are associated with infection in all conditions, a weighted gene co-expression network analysis (WGCNA) was performed (Fig 3A). This resulted in 12 to 15 different co-expression modules per temperature condition. Correlation of these modules with the infection revealed in total 12 modules (1985: four modules, 2050: three modules, Heatwave: two modules, Standard: three modules) that significantly positively correlated with the infection. These modules contained 14,341 genes, of which 1,613 genes were shared among all temperature conditions (Fig. 3B). Genes from the 12 modules were analyzed for biological function via GO term enrichment, which identified mostly translation, peptide or amide biosynthetic, peptide metabolic, and amide metabolic biological processes to be enriched (Fig. 3C). Overall, the analysis of DEGs in *U. maydis* infected plants at different temperature conditions in combination with the infection data indicates that the number of DEGs increases with increasing symptom intensity as a result of translation and peptide synthesis induction, and that the expression patterns in the cultivars become more similar.

**Fig. 3:**
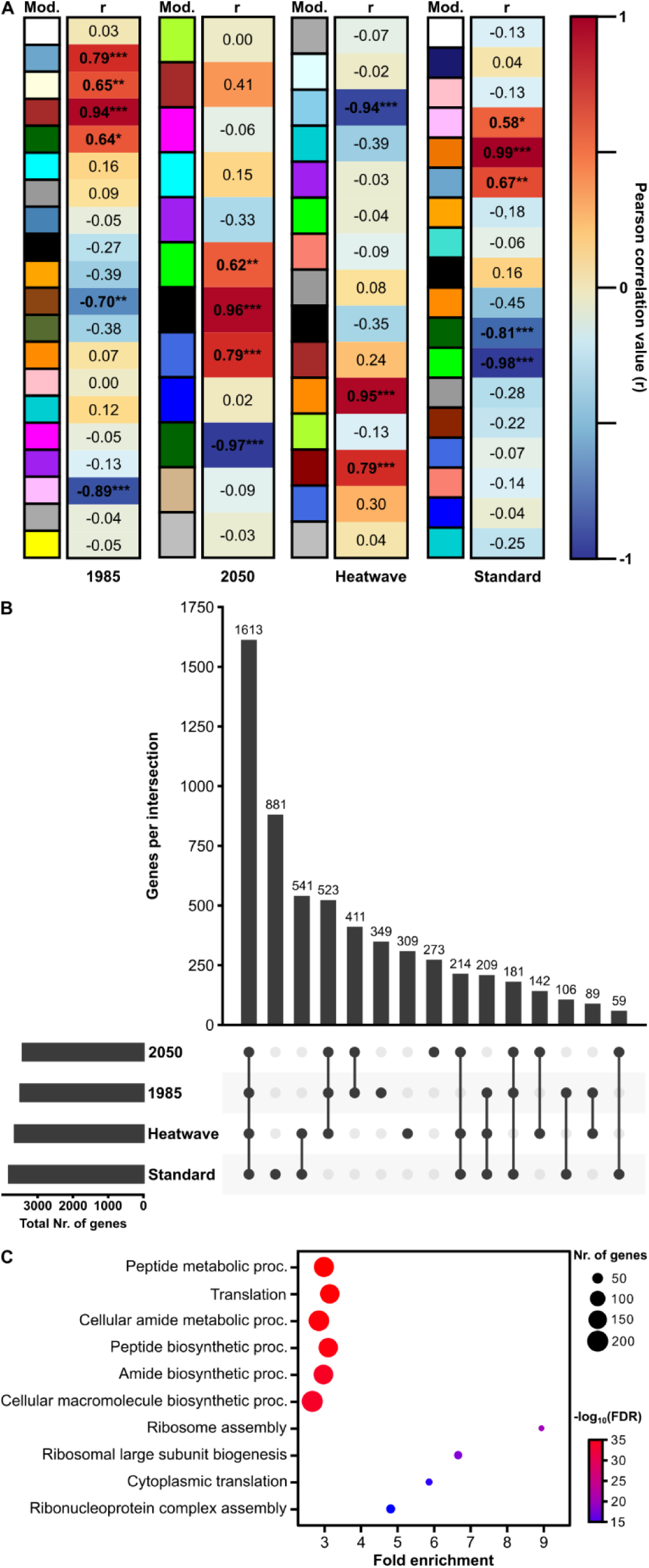
Infection-to-gene expression association under different temperatures. **A** Weighted gene co-expression network analysis module-trait correlation under different temperature conditions (temperature conditions of 1985, 2050, the temperature for 1985 with additional heatwave, or standard condition). For each temperature condition, the Pearson correlation of modules (Mod.) from the respective co-expression networks to the trait “Infection” was calculated separately. *: p-value < 0.05; **: p-value < 0.01; ***: p-value < 0.001. **B** UpSet plot of genes from weighted co-expression modules with significant positive correlation to *U. mayids* infection (r > 0.5, p-value < 0.05). For each temperature scenario the total number of genes from positively correlated modules is given. Connecting black dots indicate overlap between conditions, while vertical bars represent the number of genes per intersection. **C** Go-term enrichment analysis for biological function of genes from weighted co expression modules with significant positive correlation to *U. mayids* infection (p-value < 0.05) at all temperature conditions. The number of genes per term is indicated by dot size. FDR: false discovery rate.

### GSL4 is a resistance factor

Genes identified in the WGCNA that are strongly associated with *U. maydis* infection under all tested temperature conditions should be of general importance in the *U. maydis*-maize interaction. To test this hypothesis, we employed the Trojan Horse strategy on one small secreted maize protein identified in the WGCNA, because we found a significant enrichment of peptide-related GO terms. The Trojan Horse approach “injects” the protein of interest into the apoplast of the infected maize cell and subsequent assays can be performed, such as disease ratings, to test the impact of a protein of interest during infection ^30–32^. *GSL4* (*Gibberellic Acid Stimulated Transcript-like4*) encodes a small secreted protein with a molecular weight of 11.91 kDa and a GASA (Gibberellic Acid-Stimulated Arabidopsis) domain ^33^. Because the highest expression of *GSL4* was measured in infected dent cultivars (Figure 4A) and these cultivars have a significantly lower *U. maydis* colonization compared to flint maize (Figure 4B, C), both dent and flint cultivars were used for these experiments. In all cases, the addition of GSL4 significantly reduced symptom intensity (Fig. 4D). The greatest reduction was observed in the dent maize cultivar B73, which also had the highest *GSL4* expression tested after infection with *U. maydis* (Fig. 4A). At the same time, the relative colonization with *U. maydis* was significantly reduced in all three cultivars tested (Fig. 4E) indicating that GSL4 acts as a resistance factor.

**Fig. 4:**
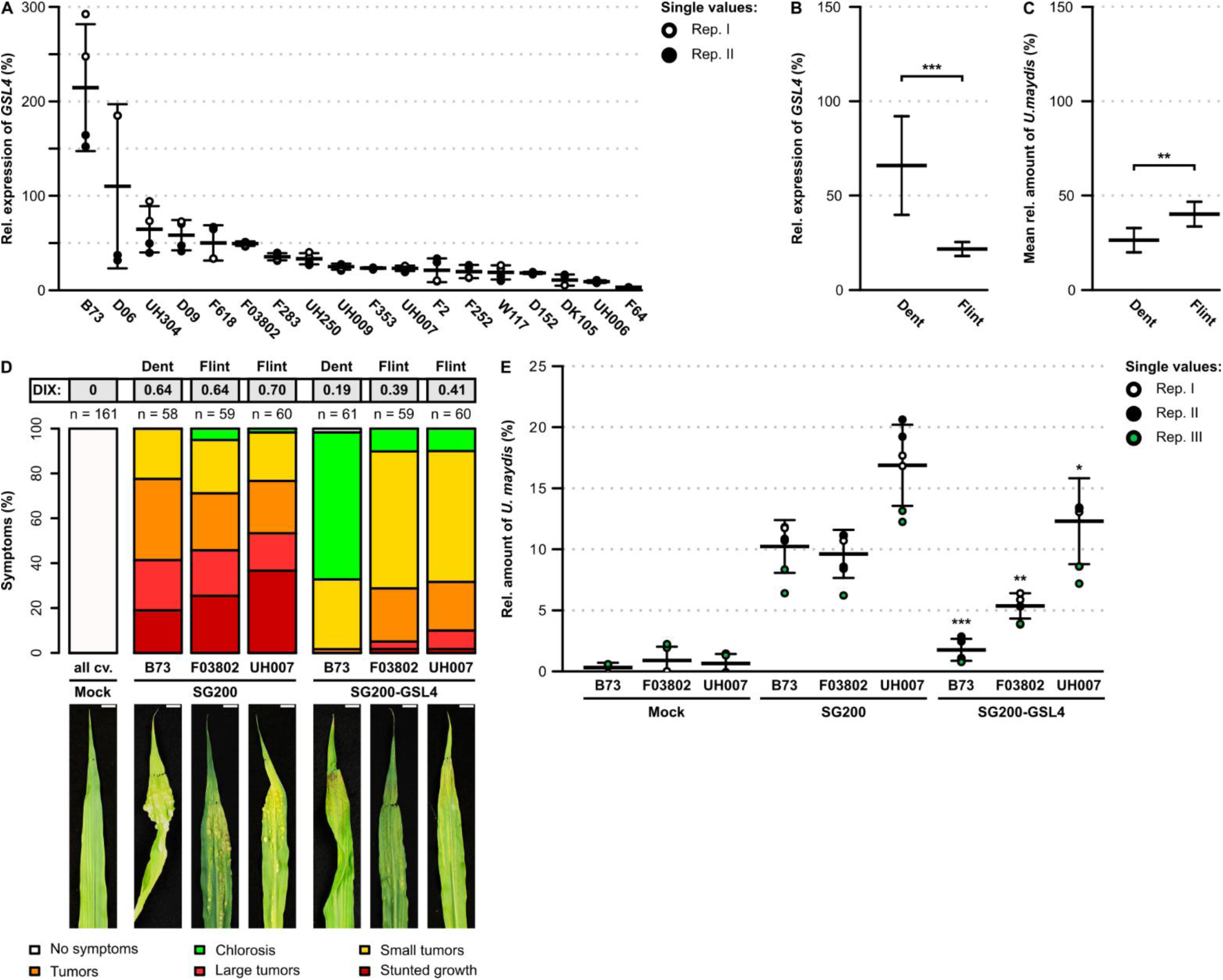
Functional analysis of GSL4 during *U. maydis* infection. **A** Relative expression of *GSL4* in 18 maize cultivars after infection at standard conditions was evaluated by qRT-PCR in comparison to *CYANASE*. qRT-PCRs were performed in two biological replicates, while each replicate consisted of two technical replicates (RepI (open circles), RepII (closed circles)). Average expression between replicates is given and error bars represent standard deviation. **B** Mean relative expression of *GSL4* and *U. maydis* biomass in dent versus flint maize cultivars. The mean relative expression of *GSL4* was calculated based on expression values of the 18 maize cultivars in (A). Relative *U. maydis* biomass was measured by qPCR in all 18 maize cultivars in two biological replicates, while each replicate consisted of two technical replicates. The mean *U. maydis* biomass of all dent or flint was calculated. Error bars indicate standard deviation. Significance was determined by Student’s *t*-test (*: p-value < 0.05; **: p-value < 0.01; ***: p-value < 0.001). **C** Disease ratings of B73 (dent), F03802 (flint), or UH007 (flint) 6 days after infection with the Trojan Horse strain SG200-GSL4, or the *U. maydis* progenitor strain SG200, or mock under standard temperature conditions. The different symptom categories are represented by specific colors. Experiments were performed in three biologically independent replicates and the total number of plants used (n) is indicated in each case. The mean infection intensity is given as disease index (DIX). Photos show representative leaves for each of the tested conditions. Scale bar: 10 mm. **D** Quantification of *U. maydis* biomass by qPCR of leaf samples. Samples were taken of B73 (dent), or F03802 (flint), or UH007 (flint) 6 days after infection with the Trojan Horse strain SG200-GSL4, or the *U. maydis* progenitor strain SG200, or mock under standard temperature conditions. qPCRs were performed in three biological replicates, while each replicate consisted of two technical replicates (RepI (open circles), RepII (closed circles), RepIII (green circles)). Average biomass between replicates is given and error bars represent standard deviation.

### GABA promotes *U. maydis* infection

In theory, samples with low *U. maydis* colonization should express higher levels of resistance factors. The lowest *U. maydis* colonization was measured at the temperature condition of 1985. Thus, a WGCNA of the 1985 RNAseq dataset was performed where the resulting modules were correlated to the relative amount of *U. maydis* colonization. Here, *U. maydis* colonization was finely dissected based on the fungal biomass data obtained by qRT-PCR. Module purple was identified, which significantly correlated to the lowest measured *U. maydis* colonization at temperature 1985 (Fig. 5A). In depth analysis of genes in this module revealed significant up-regulation of two γ-Aminobutyric acid GABA-transaminase isoforms. GABA-transaminases are involved in degradation of GABA ^34^, thus this result suggests that GABA may influence infection. Subsequently, *U. maydis*-infection experiments were performed with and without addition of GABA (Fig. 5B, C). In mock infected plants, GABA treatment had no visible effect. In contrast to that, GABA treatment during *U. maydis* infection resulted in more severe symptoms and a higher *U. maydis* biomass compared to the solvent control (H_2_O) treated plants (Fig. 5B, C). This result implies that GABA promotes *U. maydis* infection.

**Fig. 5:**
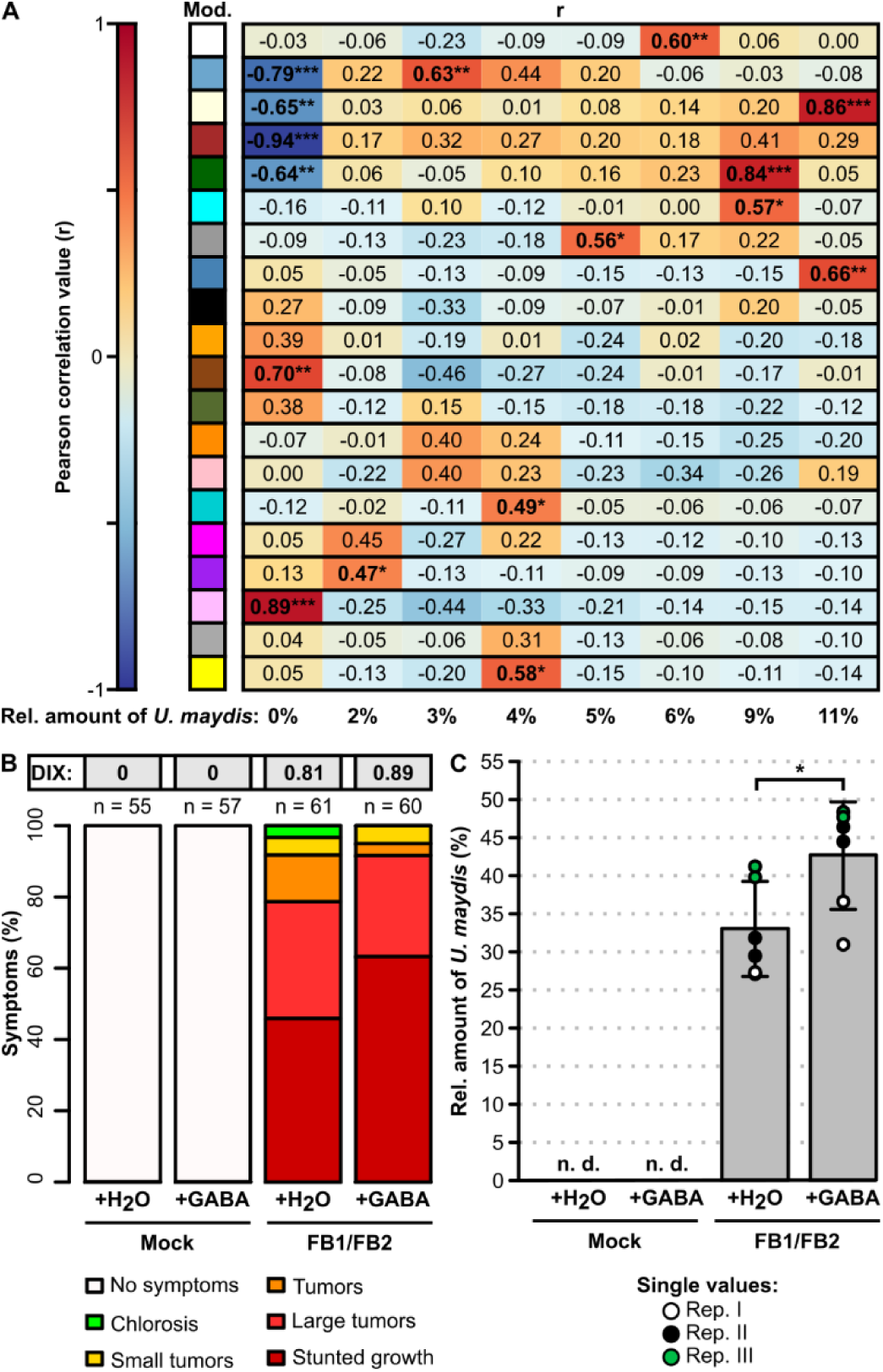
Impact of GABA on *U. maydis* infection. **A** Weighted gene co-expression module to infection severity association under the 1985 temperature condition. Co-expression modules (Mod.) were Pearson correlated to different fungal biomasses, measured by qPCR. *: p-value < 0.05; **: p-value < 0.01; ***: p-value < 0.001. **B** Influence of GABA on the *U. maydis*-maize interaction was tested by spray application of GABA on infected plants at standard temperature conditions. Mock or *U. maydis* infected plants were treated with GABA or solvent control (H_2_O) followed by disease ratings 6 dpi. The different symptom categories are represented by specific colors. Experiments were performed in three biologically independent replicates, the total number of plants used (n) is indicated in each case. The mean infection intensity is given as disease index (DIX).**C** Quantification of *U. maydis* biomass after GABA treatment by qPCR of leaf samples. Samples were taken of mock or *U. maydis* infected plants, which were treated with GABA or solvent control (H_2_O). qRT-PCRs were performed in three biological replicates, where each replicate consisted of two technical replicates (RepI (open circles), RepII (closed circles), RepIII (green circles)). Average biomass between replicates is given and error bars represent standard deviation. Significance was determined by Student’s *t*-test (*: p-value < 0.05).

## Discussion

In February 2024, news outlets around the world posted that for the first time the last 12 months were consistently 1.5 °C warmer than before industrialization ^35–37^. It is predicted that global warming will lead to agricultural yield losses of 10 - 22% by 2100 ^23,38,39^. In addition to the direct impact of rising temperatures on crops, it also is predicted that higher temperatures will lead to increased susceptibility of agricultural plants to pathogens and that pathogens will become more virulent ^24,40^. In order to test these predictions experimentally and to gain deeper insights into variations of maize cultivar specific responses to *U. maydis*, the future temperatures for the State of Bavaria (Germany) were modeled. Based on these predicted temperature changes *U. maydis*-maize infection studies were carried out on 18 different maize cultivars. In addition to disease rating data, further phenotyping data were collected and 144 RNA-seq analyses were performed.

We found that a temperature change of just 1.2 °C/1.5 °C (day/night) resulted in larger tumors, and earlier onset of melanized spore formation. Also, an increase of DEGs was measured at higher temperatures. This indicates that even small temperature increases related to climate change led to a faster development of maize smut disease. In line with this, a micro array-based study of maize seedlings infected with *U. maydis* found that the number of DEGs increases with disease progression ^5^. The higher susceptibility of maize at elevated temperatures might be partially a result of a decreased defenses response due to a compromised salicylic acid (SA) pathway as has been shown in Arabidopsis and tobacco ^41–43^. Evidently, *U. maydis* exploits different strategies to tamper with SA levels during infection ^32,44,45^. For example, the *U. maydis* effector Cmu1 (Chorismate mutases1) is translocated into the plant cell to lower the chorismate pool available for SA synthesis ^44^. Also, higher temperatures have been linked to enhanced effector secretion ^41^. Apart from that, the faster disease progression at higher temperatures might be a result of increased nutrition available to the fungus. Several studies proved that plant provided carbon sources are an important factor during infection ^3,5,46–49^. In this study an increase in maize leaf length was detected with rising temperatures indicative of higher plant biomass and photosynthetic activity. Also, it was hypothesized that differences in carbon source levels at the infection site could contribute to maize cultivar specific resistance based on observations made in the US NAM founders ^13^. Concurrently, differences in leaf length between maize cultivars after infection and variations in cultivar-dependent susceptibility occurred in this study. In the future, leaf length data in combination with the transcriptome data in this report might become a useful tool for better understanding the impact of carbon sources during infection. Also, differences in susceptibility between dent and flint maize were identified in this study. One factor responsible for these differences might be GSL4. *GSL4* expression is upregulated after infection, especially in dent maize cultivars. Trojan Horse delivered GSL4 significantly reduced symptom severity and *U. maydis* colonization indicating that GSL4 is a resistance factor. In a variety of plants over-expression of SNAKINs, which contain a GASA domain like GSL4, resulted in enhanced resistance to pathogens ^50–52^.

Our WGCNA indicated a possible influence of GABA on *U. maydis* infections. Application of GABA on infected plants resulted in enhanced symptoms and a significantly increased *U. maydis* colonization. It remains unclear whether the stronger symptoms occur because GABA is an additional nutrient source for *U. maydis* or whether GABA interferes with plant signaling pathways involved in defense. *U. maydis*-infected plants accumulate free amino acids at the infection site, and these are probably taken up by the fungus ^5,8^. Because GABA is the primary amine of glutamic acid, it cannot be excluded that GABA can be taken up and metabolized by *U. maydis*. Alternativly, GABA accumulation leads to the reduction of reactive oxygen species (ROS) ^53^. In turn, ROS are a central component of the initial plant defense response, and it has been speculated that the increased expression of antioxidant secondary metabolites during infection with *U. maydis* could be important to mitigate ROS-related cell death ^5^. Interestingly, GABA is increasingly accumulated under abiotic stress conditions, which will occur more frequently in the future due to climate change and could therefore favor *U. maydis* infections ^54^.

In summary, the phenotypic and transcriptomic datasets acquired in this study demonstrate the devastating impact of climate change-associated minimal temperature changes on plant-pathogen interactions. The increased susceptibility of maize and the faster completion of the *U. maydis* lifecycle imply an increasing risk of several infection events within one growing season accompanied by severe yield loss, which is in line with previous studies that reported that the frequency of fungal infections could be boosted and that the relative amount of fungal pathogens in the soil is increased under global warming conditions ^24,25^. Through detailed analysis of our new data, plant factors, such as GSL4 or GABA, associated with temperature changes and *U. maydis* infection could be identified in the EU NAM gene pool in the future underlining the significance of experimental data collected at temperatures reflecting climate change conditions. Then these factors then can be used in classical breeding as well as in *cis*-genetics or genome editing-mediated precision breeding to generate plants that are more adapted to the upcoming challenges of climate change.

## Methods

### Climate models

For estimations concerning future climate change, it is scientific practice to use a model ensemble of multiple climate projections. However, the projections perform very differently in reproducing the properties of the regional climate (e.g. seasonality of precipitation, regional precipitation distribution). Previously a plausibility check was used for regional climate projections in order to select a uniform data basis for statements on climate change, impact modeling and adaptation measures ^28^. As a result of this plausibility check, the Bavarian Ensemble was selected as a robust basis for analysis regarding the future climate in Bavaria for three different RCP scenarios 2.6, 4.5 and 8.5. More information about the method and the data can be found in ‘Das Bayerische Klimaprojektionsensemble - Audit und Ensemblebildung*’* ^28^.

### Plant growth conditions

Eighteen maize lines were used for the experiments: B73 as an anchor to the US Nested Associated Mapping (NAM) ^55^ panel and 17 inbred founder lines of the European NAM (EU-NAM) population (D06, D09, D152, DK105, F2, F64, F252, F283, F353, F618, F03802, UH006, UH007, UH009, UH250, UH304, W117) ^27^. Seedlings were pre-cultivated in the greenhouse as previously described ^56^. Nine days after sowing, the seedlings were moved to a walk-in climate chamber to acclimate for one day to the respective temperature condition. Here, light conditions were as follows: 25,000 lux for 14 h, one hour sunrise and sunset respectively, 8 h night.

The following temperature conditions were used: “Standard” (28°C day/25°C night), “1985” (17.9°C day/6.9°C night), “2050” (19.1°C day/8.4°C night), and “Heatwave” (35°C day/25°C night for the first 3 days after infection; 17.9°C day/6.9°C night for the remaining time of the experiment).

### *U. maydis* infections and disease ratings

Wildtype *U. maydis* strains FB1 and FB2 ^57^ were prepared as previously described ^30^. Before infection, the culture was resuspended in H_2_O to an OD_600_ of 0.3. Infection generally started at 7 p.m. For the solopathogenic *U. maydis* strains SG200 ^7^ and SG200P*pit2*::S*Ppit2*-Zm*GSL4*-*mCherry*-*HA* the cultivation procedure was similar, except that the cells were resuspended in H_2_O to an OD_600_ of 1 before infection.

The first virulence assays were performed at 6 days post infection (dpi). Depending on the temperature condition, consecutive disease ratings were performed at 10 dpi, 14 dpi, 18 dpi, and 21 dpi until spore formation was visible. Disease symptoms were scored based on the disease rating scheme proposed previously ^7^. To calculate disease indices, the numbers of plants that showed “chlorosis”, “small tumors”, “tumors”, “large tumors”, and “stunted growth” were multiplied with the factor of each category (1, 3, 5, 7, and 9, respectively) and divided by the total number of plants. The resulting value was then converted into percentage.

### Leaf length measurements

The length of the fourth leaf was measured on each day disease ratings were performed. For the measurement, the fourth leaf was stretched out on a ruler and its length was determined from the base of the sheath to the leaf blade tip. Plants with less than four leaves, a damaged fourth leaf or plants that showed a clearly delayed growth were excluded from the measurements.

### GABA treatments

For infections of GABA treated plants, 7-day-old B73-seedlings were injected with FB1/FB2. The seedlings were spray treated with 1 mM GABA (CAS: 56-12-2; Merck, Darmstadt, Germany). Spraying started on the day of infection and was repeated every two days. Per application, 20 ml of GABA-spray solution were used for 40 plants. Three independent replicates were performed at Standard temperature.

### Sample collection

Samples for gDNA and RNA extractions were collected at 6 dpi for all temperature conditions for mock or FB1/FB2 infections as described previously ^58^. Samples were collected in two independent biological replicates for all temperature conditions.

### Extraction of nucleic acids and cDNA synthesis

The collected tissue was homogenized by hand grinding in liquid nitrogen. About 200 mg of the homogenized leaf powder was used for genomic DNA (gDNA) or total RNA extractions respectively. gDNA was extracted using the MasterPure^TM^ Complete DNA & RNA Purification Kit (Biozym, Hessisch Oldendorf, Germany). Total RNA was prepared using the PureLink^TM^ RNA Mini Kit (Fisher Scientific, Schwerte, Germany). To generate cDNA, 1 µg of total RNA was reverse-transcribed with oligo(dT) primer using M-MLV Reverse Transcriptase (Thermo Fisher Scientific, Waltham, USA).

### qPCR and qRT-PCR analysis

Quantitative and quantitative real-time PCRs (qPCR and qRT-PCR) were performed on a Realplex^2^ epgradient S Mastercycler (Eppendorf) using the KAPA SYBR FAST Universal 2x Master Mix (Merck, Darmstadt, Germany) and cycling conditions described previously (van der Linde *et al.*, 2011). The specificity of the amplicon was verified via a consecutive melting curve. Generally, two technical replicates were performed for each tested sample. Primers that were used for the experiments are listed in the supp. Primer list.

To quantify the relative amount of *U. maydis* in infected leaves, the abundance of *U. maydis*-specific *ppi* and maize-specific *GAPDH* were measured by qPCR using gDNA as template ^5^. The relative amount of *ppi* to *GAPDH* was then calculated using the 2^−ΔΔct^ method ^59^. Expression of *GSL4* in infected leaves was determined by qRT-PCR and calculated relative to *CYANASE*.

### RNAseq and data analysis

Library preparation and RNAseq for nine maize cultivars (B73, D06, D09, D152, F64, F03802, UH006, UH250, W117) were performed at the Genomics Core Facility “KFB - Center of Excellence for Fluorescent Bioanalytics” (University of Regensburg, Regensburg, Germany). Sequencing was conducted on an Illumina NextSeq 2000 instrument. RNAseq data analysis was performed using R (https://www.r-project.org/) in an Ubuntu-based Jupyter Notebook (https://jupyter.org/). For each sample, around 30 million reads were generated by Illumina sequencing. The quality of the raw reads was checked using FastQC (v0.12.1) ^60^ and MultiQC (v1.14) ^61^. Adapters were trimmed using the Trimmomatic software (v0.39) ^62^, which also removed reads below 36 nucleotides (Survival rate: 97-99%). Trimmed reads were mapped to the B73 *Z. mays* reference genome version 5 via HISAT2 (v2.2.1) ^63^ and quantified using featureCounts (v2.0.1) ^64^. For mock as well as *U. maydis* infected samples, about 30 to 50 million reads were counted at each temperature condition. The approximately 40,000 identified genes per sample were filtered to remove genes with counts below 10, leaving around 27,000 genes per sample. The prefiltered genes were then normalized and subjected to differential gene expression via DESeq2 (v1.38.3) ^65^, testing *U. maydis*-infected samples versus mock. This was done for each cultivar at each temperature condition separately to reduce variation. Genes with a log_2_-fold change (LFC) > 2 and Benjamini–Hochberg-adjusted P value < 0.001 were considered significantly upregulated, genes with a LFC < 2 and Benjamini–Hochberg-adjusted P value < 0.001 were considered significantly downregulated. Unique differentially expressed genes (DEGs) of all lines were merged for each temperature condition to generate a complete list of all DEGs at a certain temperature. Line-specific (DEGs present only in single maize cultivars), shared (DEGs present in at least two maize cultivars), and core DEGs (DEGs present in all analyzed maize cultivars) were determined by comparing DEGs of single cultivars to the DEGs of remaining cultivars. To construct co-expression networks, the WGCNA package (v1.72) ^66^ was used. The 10,000 most variable genes which were selected from filtered counts (>10 counts) of mock and FB1/FB2 infected samples were transformed into FPKM (fragments per kilobase per million mapped fragments) for each temperature condition separately and used as input for co-expression analysis. The samples were first clustered by hierarchical clustering (method = “average”) and the best power to reach scale free topology was determined for each temperature condition (1985: R^2^ = 0.885, 2050: R^2^ = 0.844, Heatwave: R^2^ = 0.844, Standard: R^2^ = 0.819). Then, signed, Pearson-correlated co-expression networks were calculated using the function exp2gcn ^67^ with a module merging threshold of 0.8. For each module, module size was determined and the module eigengene expression was calculated. The eigengene expression was then Pearson correlated to the temperature condition, the disease index, and the relative amount of *U. maydis* respectively. Genes from correlating modules were subjected to a GO term enrichment analysis (p-value cutoff: 0.05) using ShinyGO ^68^.

### Plasmid construction and *U. maydis* transformation

The full-length sequence of *GSL4* was cloned from cDNA of B73. The product was cloned into the p123-P_pit2_-*pit2*-*mCherry*-*Ha* vector backbone ^69^. The resulting plasmid was then transformed into the *U. maydis* strain SG200 and its functionality was verified as described previously ^30^.

## Acknowledgement

The authors would like to thank the Chair of Plant Breeding at TUM (Germany), the INRA (France), the CIAM (Spain), and the University of Hohenheim (Germany) for providing EU NAM founder cultivars. Funding was provided by the Deutsche Forschungsgemeinschaft (DFG) SFB924 Project A14 and the Bayerisches Staatsministerium für Umwelt und Verbraucherschutz (Bavarian State Ministry of Environment and Consumer Protection) BayKlimaFit 2 Project TP7.

## Author contributions

C.S. and K.v.d.L. designed experiments, conducted experiments, performed data analysis, and wrote the manuscript. F.H. and C.Z. designed experiments, conducted experiments, and performed data analysis.

## Data availability

Source data are provided in this paper. All data supporting the findings of this study that are not directly available within the paper (and its supplementary data) will be upon reasonable request available from the corresponding authors (KvdL). The RNA sequencing data have been submitted to the NCBI Sequence Read Archive (SRA) (accession number: XXX).

